# Preexisting mycobacterial infection modulates *Candida albicans*-driven pyroptosis

**DOI:** 10.1101/2021.08.04.455086

**Authors:** Bharat Bhatt, Praveen Prakhar, Gaurav Kumar Lohia, Raju S. Rajmani, Kithiganahalli Narayanaswamy Balaji

## Abstract

Active tuberculosis patients are at high risk of co-infection with opportunistic fungal pathogen *C. albicans*. However, the molecular mechanisms that orchestrate pathogenesis of *Mycobacterium tuberculosis* (Mtb)-*C. albicans* co-infection remains elusive. In the current study, we utilise a mouse model to demonstrate that Mtb promotes macrophage environment conducive for *C. albicans* survival. Mtb-dependent PKCζ-WNT signalling axis induces expression of an E3 ubiquitin ligase, COP1. A secondary infection of *C. albicans* in such Mtb-infected macrophages causes COP1 to mediate the proteasomal degradation of IRF9, a cardinal factor that we identified to arbitrate an inflammatory programmed cell death, pyroptosis. *In vivo* experiments mimicking a preexisting Mtb infection demonstrate that inhibition of pyroptosis in mice results in increased *C. albicans* burden and aberrant lung tissue architecture, leading to increased host mortality. Together, our study reveals the crucial role of pyroptosis regulation for manifesting a successful *C. albicans*-Mtb co-infection.

## Introduction

*Candida albicans* is an opportunistic pathogen, associated with high mortality rate of up to 40-58% in immunocompromised patients [1]. Susceptibility to *C. albicans* infection is further compounded by factors ranging from prior viral or bacterial infection to various antibiotic treatments that weakens the host immune system. In this context, there exists a number of clinical studies that report up till 58% incidence of *Candida* infection in active tuberculosis patients [2–5]. *Mycobacterium tuberculosis* (Mtb), a causative agent of tuberculosis, is one of the leading cause of death worldwide [6]. Mtb thrives in the host by creating permissible environment for its own survival by skewing various host responses including cytokine response towards the anti-inflammatory response, inhibiting apoptosis and autophagy [7–10]. Further, stringent and long course of anti-tubercular treatment makes the patient more susceptible to opportunistic fungal infections [11]. Therefore, it becomes elemental to apprehend host defences against *C. albicans* during *C. albicans*-Mtb co-infection.

*C. albicans* harbours an arsenal of virulent factors amongst which transition from yeast to hyphae form is one of the key virulent factor, with yeast form predominantly existing as commensal and hyphae as invasive pathogenic form [12,13]. Phagocytosis of *C. albicans* culminates in death of macrophages and till recently, physical rupture of cell membrane by its hyphal form was considered to be the primary cause of host cell death. However, recent reports suggest that death of macrophages occurs in two distinct phases: Early phase (6 h to 8 h post infection) and late phase (10 h to 24 h post infection). Moreover, host responses elicited by *C. albicans* in early phase are very different from late phase of macrophage death. In this perspective, *C. albicans* triggered pyroptosis, is considered to be a major contributor of macrophage killing during early phase of infection [14,15].

Pyroptosis is an inflammatory, programmed cell death marked by cell rupture and concomitant increased secretion of inflammatory cytokines such as IL-1β and IL-18 in extracellular milieu. During pyroptotic cell death, inflammasome formation leads to activation of Caspase-1 or Caspase-11, leading to cleavage of Gasdermin D (GSDMD) along with cleavage of pro-IL-1β and pro-IL-18 into active IL-1β and active IL-18. Cleaved N-terminal domain of GSDMD perforates the cell membrane which actuates osmotic lysis of cell and release of IL-1β and IL-18 [16,17]. During *C. albicans* infection, NLRP3 inflammasome mediated activation of Caspase-1 has been demonstrated to play vital role in induction of pyroptosis [15]. On the contrary, execution of cell death upon Mtb infection is independent of Capase-1/Caspase-11 [18,19], hence argued against the role for pyroptosis during Mtb infection. Nonetheless, a recent literature provide differing views on Mtb role in GSDMD-mediated pyroptosis [20]. Moreover, it has been shown that Rv3364c, an Mtb protein, inhibits activation of Caspase-1 [21]. With this premise, we explored the role for pyroptosis during *C. albicans* and Mtb co-infection scenario.

Here, we report for the first time that Mtb significantly reduces the early pyroptotic macrophage death induced by *C. albicans*. Further, we found that *C. albicans* infection markedly upregulates the expression of IRF9 (Interferon Regulatory Factor 9) which leads to activation of GSDMD driven pyroptosis. More importantly, prior Mtb infection targets IRF9 for proteasomal degradation by an E3 ubiquitin ligase, COP1 (Constitutive photomorphogenesis protein 1). We found that PKCζ is crucial for the activation of WNT signalling pathway upon Mtb infection and activation of PKCζ-WNT signalling axis in turn induces the expression of COP1. Finally, *in vivo* studies demonstrate that mice co-infected with Mtb and *C. albicans* have exacerbated pulmonary pathology, decreased survival and increased *C. albicans* burden. Altogether, our results demonstrate that pyroptosis is a crucial host defence against *C. albicans* infection which is compromised upon prior Mtb infection.

## Material and Methods

### Cells and Mice

RAW 264.7 mouse macrophage cell line (obtained from National Centre for Cell Sciences, Pune, India) and peritoneal macrophages isolated from wild type BALB/c maintained at the Central Animal Facility, Indian Institute of Science were cultured in DMEM (GIBCO, Invitrogen Corporation, USA) supplemented with 10% heat inactivated FBS (Sigma-Aldrich, USA), maintained at 37°C in 5% CO_2_. Experiments with mouse macrophages were carried out after the approval from the Institutional Ethics Committee for animal experimentation as well as from Institutional Biosafety Committee. Peritoneal macrophages were isolated by intraperitoneally injecting 1ml of 8% Brewer thioglycolate medium and post 4 days, peritoneal exudates were harvested in 1X PBS.

### Bacteria and Fungus

Mid log phase culture of Mtb H37Rv grown in Middlebrook 7H9 medium (Difco, USA) supplemented with 10% oleic acid, bovine serum albumin, dextrose, catalase (OADC) were used for all experiments. Single cell suspensions of Mtb H37Rv were prepared by serially passing the culture from 24G, 26G and 30G syringe and used at an MOI of 1:10 unless indicated otherwise. Mtb H37Rv was a kind research gifts from Dr. Kanury V.S. Rao, ICGEB, India. tdTomato Mtb H37Rv was a kind research gift from Dr. Amit Singh, IISc, India. All studies involving Mtb H37Rv were carried out at the BSL-3 facility at Centre for Infectious Disease Research (CIDR), IISc. Wild-type *C. albicans* (SC5314) was procured from Microbial Type Culture Collection, Institute of Microbial Technology, Chandigarh, India (MTCC 4748). GFP expressing strain of *C. albicans* (CEC2684) was a kind research gift from Dr. Kaustuv Sanyal, JNCASR, India. *C. albicans* were grown to mid log phase in YPD medium at 28°C. Culture was washed in 1XPBS and *in vitro* infection was performed at an MOI of 4:1.

### Reagents and antibodies

General laboratory chemicals were purchased from Sigma-Aldrich (USA) or Promega (USA) or Himedia (India) or Merck (Germany) for all the experiments. Tissue culture plastic ware was obtained from BD Falcon (USA) or Tarsons (India). Cell culture antibiotics were purchased from Sigma-Aldrich (USA). Anti-Caspase-1 and anti-K48-linkage specific polyubiquitin, phospho-GSK3β and phospho-β-CATENIN antibodies were purchased from Cell Signaling Technology (USA). Anti-COP1 antibody was procured from Santa Cruz Biotechnology (USA), anti-IRF9 and anti-GSDMD from Abcam, Anti-β-actin (AC-15) antibody was obtained from Sigma-Aldrich (USA) and HRP conjugated anti-rabbit IgG antibody was obtained from Jackson ImmunoResearch (USA). 4′,6-Diamidino-2-phenylindole dihydrochloride (DAPI) and chloramphenicol was from Sigma-Aldrich. Haematoxylin and Eosin stain solution was from Thomas Baker.

### Treatment with pharmacological reagents

For all *in vitro* experiment, inhibitor treatment was given 1h prior to infection: Ac-YVAD-CMK (50mM) was purchased from Cayman Chemicals. IWP2 (5μM), β-CATENIN inhibitor (15μM), PKCζ inhibitor (5μM) and MG-132 (20μM) were purchased from Calbiochem (USA). MG-132 treatment was given 4h prior to completion of experiment. 0.1% DMSO was used as the vehicle control.

### RNA isolation and quantitative real-time PCR

Total RNA was isolated from peritoneal macrophage cells or THP1 cells or RAW 264.7 cells using TRI Reagent (Sigma-Aldrich, USA) as per the manufacturer’s protocol. For RT-PCR, using First strand cDNA synthesis kit (Bioline, UK), 1 μg of total RNA was converted into cDNA. SYBR Premix Ex Taq II (Takara) was used for quantitative real time PCR for quantification of target gene expression. Experiments were repeated at least three times independently to confirm the reproducibility of the results. *Gapdh* amplification was used as internal control. Primer sequences are listed in Supplementary Table 1. Primers were synthesized and obtained from Eurofins Genomics Pvt. Ltd. (India).

### Preparation of cell lysate and Immunoblotting

Immunoblotting was performed using whole cell lysate of cells to check protein levels. Cells were washed with PBS. Cell were lysed in RIPA buffer [50 mM Tris-HCl pH 7.4, 1% NP-40, 0.25% sodium-deoxycholate, 150 mM NaCl, 1 mM EDTA, 1 mM PMSF, 1 mM Aprotinin, 1 mM Leupeptin, 1 mM Pepstatin, 1 mM Na3VO4, 1 mM NaF] and incubated for 30 min on ice. Lysed cells were then centrifuged at 13000 rpm for 10 min at 4°C to obtain supernatant for whole cell protein lysate. Protein concentrations were determined by Bradford’s method. Equal protein amount were used to perform SDS-PAGE and transferred onto polyvinylidene difluoride membranes (PVDF) (Millipore, USA) by the semi-dry transfer (Bio-Rad, Australia) method. Nonspecific binding was blocked with 5% non-fat dry milk powder in TBST [20 mM Tris-HCl (pH 7.4), 137 mM NaCl, and 0.1% Tween 20] for 1h. Blots were then incubated at 4°C overnight with primary antibodies in 5% BSA (in TBST). After washing in TBST, blots were incubated with goat anti-rabbit IgG secondary Antibody conjugated to HRP in 5% BSA for 2h. After washing in TBST, the blots were developed with the ECL system (Perkin Elmer, USA) as per manufacturer’s instructions.

### Transient transfections

1 million RAW 264.7 cells were transfected with PKCζ dominant negative construct using PEI reagent. After 48h of transfection, cells were treated or infected as indicated. For siRNA transfection, mouse peritoneal macrophages were treated with 100μM of specific siGENOME SMARTpool siRNA (Dharmacon) using PEI reagent. After 36h of transfection, cells were treated as indicated.

### Immunofluorescence (IF)

Macrophages were fixed with 3.6% formaldehyde for 30 min at room temperature. The cells were washed with PBS and 2% BSA in PBST was used for blocking. After blocking, cells were stained with GSDMD at 4°C overnight. Following overnight staining, cells were incubated with Alexa488-conjugated secondary antibody for 2h and nuclei were stained with DAPI. The coverslips were mounted on a slide with glycerol. In case of cryosection, frozen sections were allowed to thaw at room temperature. Following blocking using 2% BSA containing 0.02% saponin, specific antibodies were used to stain the sections at 4°C overnight. Sections were then incubated with Alexa488-conjugated secondary antibody for 2 h and DAPI was used to stain nuclei. Sections were mounted on by coverslip with glycerol as the medium. Zeiss LSM 710 Meta confocal laser scanning microscope (Carl Zeiss AG, Germany) was used to capture confocal images and images were analysed using ZEN 2009 software. Corrected Total Cell Fluorescence (CTCF) was calculated using the following formula: CTCF = (Area of selected cell X Mean florescence of cell) – (Area of selected cell X Mean florescence of background reading).

### PI staining for viability

After completion of experiment, cells were washed with PBS followed by staining with 10μg/ml of PI (Sigma-Aldrich (USA)) for 15 min. Cells were then washed thrice with PBS followed by fixation with 3.7% PFA for 30 min. Nuclei were stained with DAPI.

### Enzyme linked Immunosorbent Assay (ELISA)

Cell culture supernatant was harvested and used to carry out ELISA for IL-1β (Invitrogen) and IL-18 (Invitrogen) as per the manufacturer’s instruction. Briefly, 96-well flat bottom plate (Nunc MaxiSorp, Thermo Scientific) was coated with specific capture antibody at 4°C overnight. Following this, wells were washed with 1X PBST (1X PBS with 0.05% Tween 20) and blocked with blocking buffer for 2 h at room temperature. Following two washes with wash buffer, 50μl of Sample Diluent C was added along with 50μl of culture supernatant. To above, 50μl of Detection antibody was added and was incubated at room temperature for 2 h. Wells were washed with wash buffer and were incubated with specific streptavidin/avidin-HRP antibody for 1 h. Following this, wells were washed and substrate solution (tetramethylbenzidine; TMB) was added. Reaction was arrested using 2N H_2_SO_4_ after 15min and absorbance was measured at 450 nm and 570nm using an ELISA reader (Tecan, Switzerland).

### Hematoxylin and Eosin (H&E) staining

Upper right lung lobes were fixed in formalin, embedded in paraffin and microtome sections of 5 μm thickness were obtained using microtome. Microtome sections were then subjected to deparaffinization followed by rehydration. As per the manufacturer’s protocol, sections were stained with haematoxylin followed by eosin staining. Then sections were dehydrated, and were mounted with coverslip using D.P.X. mountant (Fisher Scientific).

### Cryosection preparation

The excised fixed lungs were placed in the optimal cutting temperature (OCT) media (Jung, Leica). Using Leica CM 1510 S or Leica CM 3050 S cryostat, cryosections of 10 μm were prepared with the tissue embedded in OCT being sectioned onto glass slides and then stored at −80°C.

### Immunoprecipitation assay

Post treatment, cells or lung tissues were washed with ice-cold PBS and gentle lysis with ice-cold RIPA buffer was performed. Cell or lung lysate was then incubated with anti-COP1, anti-IRF9, anti-UbK48 or Rabbit IgG at 4°C for 2 h on cyclomixer followed by 4h incubation with pre-blocked protein A beads (Bangalore Genei). Beads were then harvested and washed and boiled in 5X Laemmli buffer for 10 min. The samples were separated by SDS-PAGE and immunoblotting for indicated protein was performed.

### Chromatin immunoprecipitation (ChIP) assay

Macrophages were fixed with 3.7% formaldehyde for 15 min at room temperature followed by addition of 125 mM glycine to inactivate formaldehyde. 0.1% SDS lysis buffer containing 50 mM Tris-HCl (pH8.0), 200 mM NaCl, 10 mM HEPES (pH 6.5), 0.1% SDS, 10 mM EDTA, 0.5 mM EGTA, 1 μg/ml of each aprotinin, leupeptin, pepstatin, 1 mM Na3VO4 and 1 mM NaF was used to lyse nuclei. Bioruptor Plus (Diagenode) was used at high power for 70 rounds of 30 sec pulse ON/45 sec OFF to shear the chromatin. Chromatin extracts harbouring DNA fragments with an average size of 500 bp were immunoprecipitated using specific antibody and anti-rabbit IgG complexed with Protein A agarose beads (Bangalore Genei). Immunoprecipitated complexes were then sequentially washed [Wash Buffer A: 50 mM Tris-HCl (pH8.0), 500 mM NaCl, 1 mM EDTA, 1% Triton X-100, 0.1% Sodium deoxycholate, 0.1% SDS and protease/phosphatase inhibitors; Wash Buffer B: 50 mM Tris-HCl (pH 8.0), 1 mM EDTA, 250mM LiCl, 0.5% NP-40, 0.5% Sodium deoxycholate and protease/phosphatase inhibitors; TE: 10 mM Tris-HCl (pH 8.0), 1 mM EDTA] and eluted in elution buffer [1% SDS, 0.1 M NaHCO3]. Eluted samples were then treated with RNase A and Proteinase K, DNA was precipitated using phenol-chloroform-ethanol method. Quantitative real time PCR was used to analyse purified DNA. Values obtained in the test samples were normalized to amplification of the specific gene in Input and IgG pull down and represented as fold enrichment. Primer sequences are listed in Supplementary Table 2.

### Mouse model of Co-infection

BALB/c mice were aerosolized with 200 CFU of Mtb in an aerosol chamber (Wisconsin-Madison, USA) and CFUs were confirmed by plating the homogenized lung tissue on Middlebrook 7H10 (Difco) agar. Post 28 days, intravenous infection of 1*10^6^ *C. albicans* was given via tail vein injection using previously described protocol for systemic Candidiasis [22]. Survival of mice was assessed. For fungal burden, mice were sacrificed, spleen and left lung lobe were homogenized in sterile PBS, serially diluted and plated on YPD agar containing chloramphenicol (50μg/ml).

### *In vivo* mouse model for inhibitor treatment

BALB/c mice were intraperitoneally administered with Ac-yvad-cmk (12.5μmol/kg) for 2 days at an interval of 24h. Post 12h of 2^nd^ dose of Ac-yvad-cmk, intravenous infection of 1*10^6^ *C. albicans* was given via tail vein injection. Survival of mice, fungal burden and lung pathology were studied as described above.

### Statistical analysis

Student’s t-test distribution and one-way ANOVA or two-way ANOVA followed by Tukey’s multiple-comparisons were used to determine the levels of significance for comparison between samples. The data in the graphs are depicted as the mean ± S.E for the values obtained from at least 3 or more independent experiments and P values < 0.05 were defined as significant. Log-rank (Mantel-Cox) test was used for mice survival experiments. For all the statistical analysis GraphPad Prism 5.0 software was used.

## Results

### *M. tuberculosis* co-infection enhance severity of *C. albicans* pathogenesis

To understand the interplay between these two pathogens, we established an *in vivo* model of co-infection wherein BALB/c mice were aerosolised with Mtb at ~200 CFU leading to granuloma formation in 4 weeks which marks the onset of active tuberculosis [9,23], followed by intravenous infection of *C. albicans* leading to systemic candidiasis [22]. Evidently, the mice that were co-infected showed higher mortality rate as compared to that of only *C. albicans* infected, though no mortality was observed in only Mtb infected mice for indicated time (Fig 1A). Further, burden of *C. albicans* in lung and spleen, wherein both pathogens are reported to colonize, was found to be higher during co-infection as compared to *C. albicans* infection alone (Fig 1B). Histopathology analysis of lung sections harvested from Mtb infected mice demonstrated granuloma formation with dense aggregates of lymphocytes and histiocytes. Three to four granulomas were observed at the periphery of lung while six to seven scattered foci in alveolar septa in rest of the lung parenchyma were also observed. Mice granuloma do not show necrotic caseum. In case of *C. albicans* infection, focal aggregates of lymphocytes and histiocytes were found in alveolar septa and alveolar spaces were filled with edema. A notable increase in fluid filled alveoli (pulmonary edema) was observed in lungs of mice co-infected with Mtb and *C. albicans* as compared to only *C. albicans* infected mice. Moreover, focal septal aggregates of macrophages and few lymphocytes were observed (Fig 1C), indicating distorted lung architecture and severe lung pathology during co-infection compared to Mtb or *C. albicans* infection alone.

**Fig. 1.**
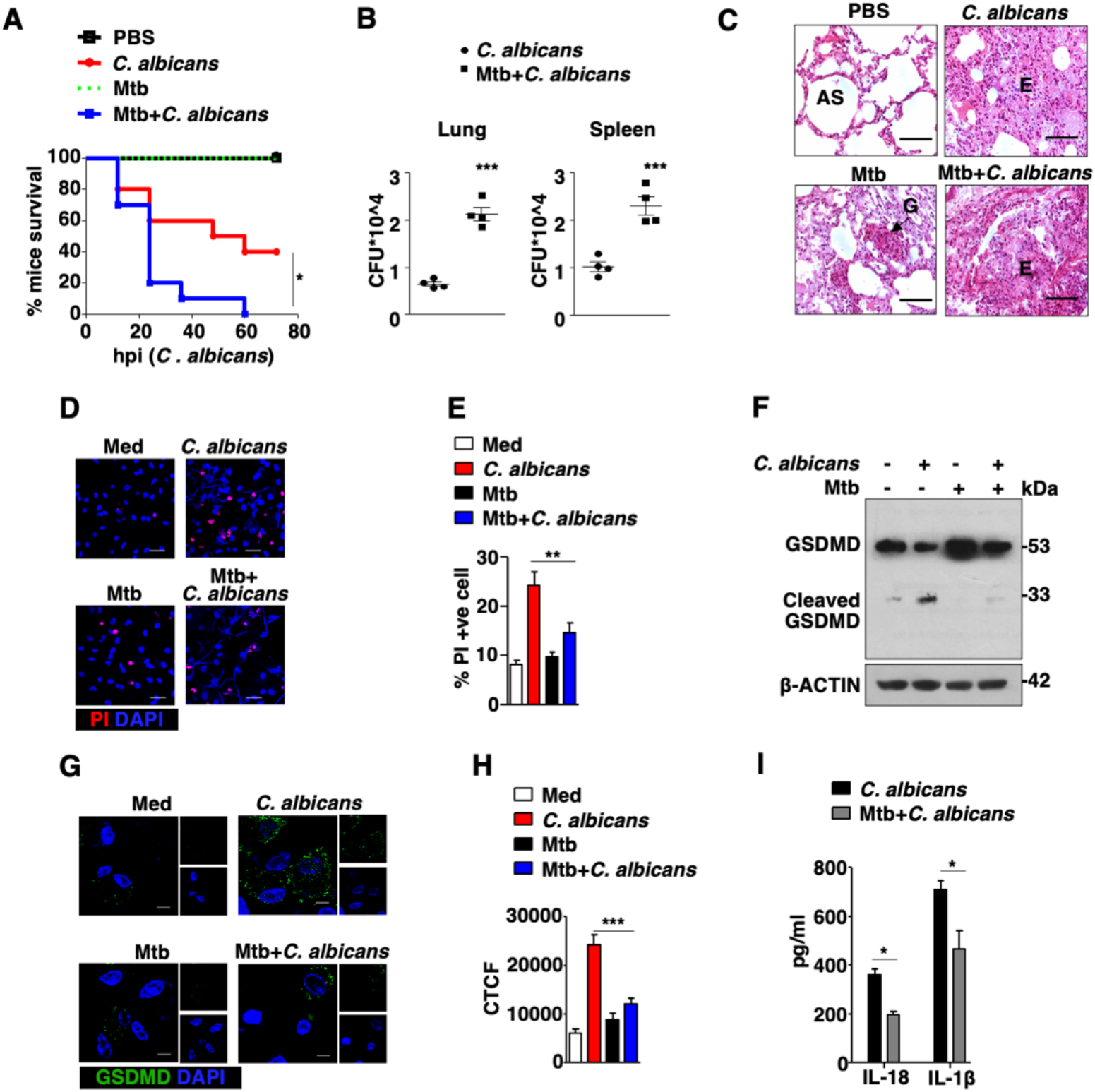
*M. tuberculosis* co-infection enhances severity of *C. albicans* pathogenesis. (A-C) Mice were infected with *C. albicans* and/or *M. tuberculosis*. (A) Survival of mice from each group (n=10). Post 24 h of intravenous infection with *C. albicans*, lungs and spleen were harvested. (B) Burden of *C. albicans* in lung and spleen, and (C) representative image of hematoxylin and eosin (H&E) stained lung sections (40X photomicrographs). (D-I) Peritoneal macrophages were infected with *C. albicans* for 6 h or Mtb for 18 h or co-infected with Mtb for 12 h followed by *C. albicans* for 6 h. (D) Representative images and (E) quantification of peritoneal macrophages stained with PI and DAPI for viability analysis. (F) Immunoblot to assess cleavage of GSDMD. (G) Representative IF images and its quantification (H) to assess surface localisation of GSDMD was performed. (I) ELISA for IL-18 and IL-1β was performed in cell culture supernatant. Error bar represents the mean ± SEM for at least 3 independent experiments; *, p < 0.05; **, p < 0.01; ***, p < 0.001 (Log-rank (Mantel-Cox) test for panel A, Student’s t-test for panel B, One way ANOVA followed by Tukey’s test for panel E and H), all blots are representative of 3 independent experiments. Scale bar, 200 μm(C), 20 μm(D) and 5 μm(G); original magnifications 40X(D) and 63X(G); Med, medium; CTCF, corrected total cell fluorescence; hpi, hours post infection; AS, alveolar space; E, eosinophils; G, granuloma.

Significantly, uptake of *C. albicans* in macrophages is independent of Mtb infection as shown in Fig S1. However, death of *C. albicans* infected macrophages was reduced upon prior infection with Mtb (Fig 1D-E). Death of macrophage occurred as early as 6 h post *C. albicans* infection and pyroptosis has been reported to be the major contributor of cell death during the early phase of infection. In this regard, cleavage and surface localisation of GSDMD (a key effector of pyroptosis) was validated to assess the level of pyroptosis. As shown, cleavage and surface localisation of GSDMD was increased upon *C. albicans* infection alone, however, to our surprise prior Mtb infection strongly reduced *C. albicans* triggered cleavage and surface localisation of GSDMD (Fig 1F-H). Further, pyroptosis is marked by increased secretion of IL-18 and IL-1. In line with GSDMD cleavage and surface localisation, secretion of IL-18 and Il-1 β was also found compromised upon prior Mtb infection.

### Caspase-1 mediated pyroptosis protects mice against *C. albicans* infection

Cleavage of GSDMD is regulated by Caspase-1 activation and in this context inhibition of Caspase-1 using Ac-yvad-cmk (selective and irreversible Caspase-1 inhibitor) upon *C. albicans* infection rescued *C. albicans* mediated cell death (Fig 2A-B, Fig S2A) along with decreased GSDMD cleavage and its surface localisation (Fig 2C-E), which is in line with literature suggesting *C. albicans* induced pyroptosis is predominantly dependent on Caspase-1 activation [14,15].

**Fig. 2.**
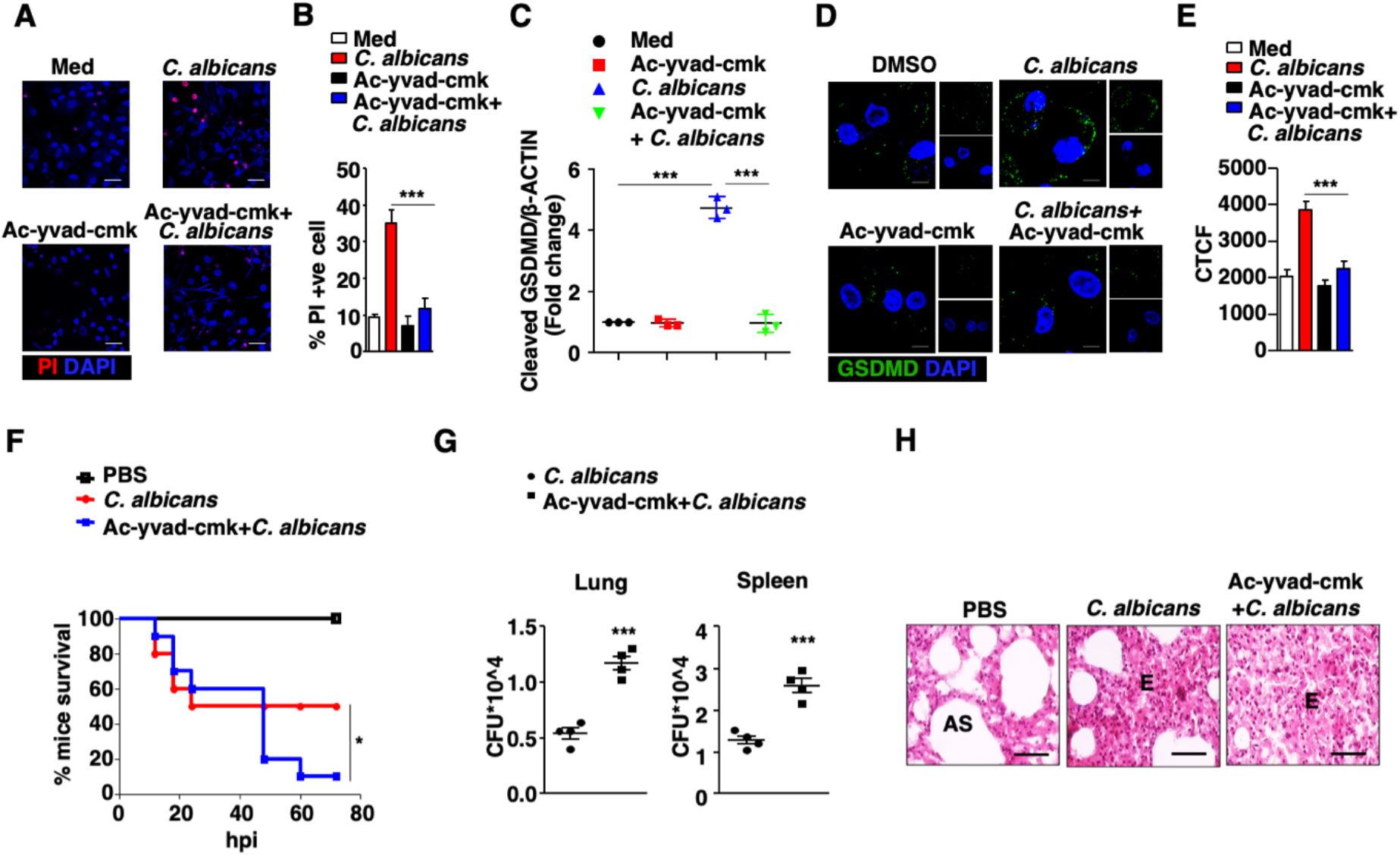
Caspase-1 mediated pyroptosis protects mice against *C. albicans* infection. (A-E) Peritoneal macrophages were infected with *C. albicans* for 6 h with or without treatment with Ac-yvad-cmk (Caspase-1 inhibitor) 1 h prior to *C. albicans* infection. Cell viability assessed by PI and DAPI staining, (A) representative fluorescence images and (B) quantification of PI positive macrophages. (C) Densitometry analysis of cleaved GSDMD. (D) Representative IF images of GSDMD and (E) its quantification. (F-H) Mice were treated with Ac-yvad-cmk followed by intravenous infection with *C. albicans*. (F) Survival of mice from each group (n=10). 24 h post infection, (G) burden of *C. albicans* in lung and spleen, (H) representatives of H&E stained lung section to evaluate lung pathology (40X photomicrographs). Error bar represents the mean ± SEM for at least 3 independent experiments; *, p < 0.05; ***, p < 0.001 (One-way ANOVA followed by Tukey’s test for panel B and E, Log-rank (Mantel-Cox) test for panel F, Student’s t-test for panel G). Each blot is representative of 3 independent experiments. Scale bar, 20 μm(A), 5 μm(D) and 200 μm(H); original magnifications 40X (A) and 63X (D); Med, medium; CTCF, corrected total cell fluorescence; hpi, hours post infection; AS, alveolar space; E, eosinophils.

In order to study *in vivo* relevance of pyroptosis, *C. albicans* infected mice were treated with Ac-yvad-cmk which led to an evident inhibition of Caspase-1 activity in lung tissue of mice (Fig S2B). We observed an increased mortality of mice treated with Ac-yvad-cmk compared to untreated mice (Fig 2F). Moreover, *C. albicans* burden in lung and spleen was higher in Caspase-1 inhibitor treated mice (Fig 2G). Accordingly, lung sections of Caspase-1 inhibitor treated mice showed concomitant infiltration of immune cells especially macrophages and eosinophils along with significant increment in edema filled alveoli and loss of alveolar spaces (Fig 2H). Together, observations from Fig 1 and Fig 2 suggest the beneficial role of pyroptosis in countering *C. albicans* pathogenesis and its restriction upon prior Mtb infection.

### IRF9 mediates *C. albicans* induced pyroptosis

Interferon Regulatory Factor (IRF) family of proteins play important role in pyroptosis as reported for various infections. For examples, IRF1 is reported to promote pyroptosis during LPS induced lung injury while IRF2 transcriptionally regulate expression of GSDMD. Similarly, IRF8 regulates pyroptosis during *Salmonella typhimurium*, *Burkholderia thailandensis*, or *Pseudomonas aeruginosa* infection [24–26]. IRF7 physically interacts with GSDMD to promote pyroptosis [27]. With this premise, we assessed the transcript levels of all the members of IRF family and found that IRF3, IRF7 and IRF9 were significantly upregulated as early as 4 hours post *C. albicans* infection (Fig 3A). Various studies have reported conflicting findings in regard to role for IRF3 or IRF7 in mediating pyroptosis [27,28]. Interestingly, IRF9 is known to transcriptionally regulate the expression of IRF7[29], yet the role for IRF9 during pyroptosis has not been explored.

**Fig. 3.**
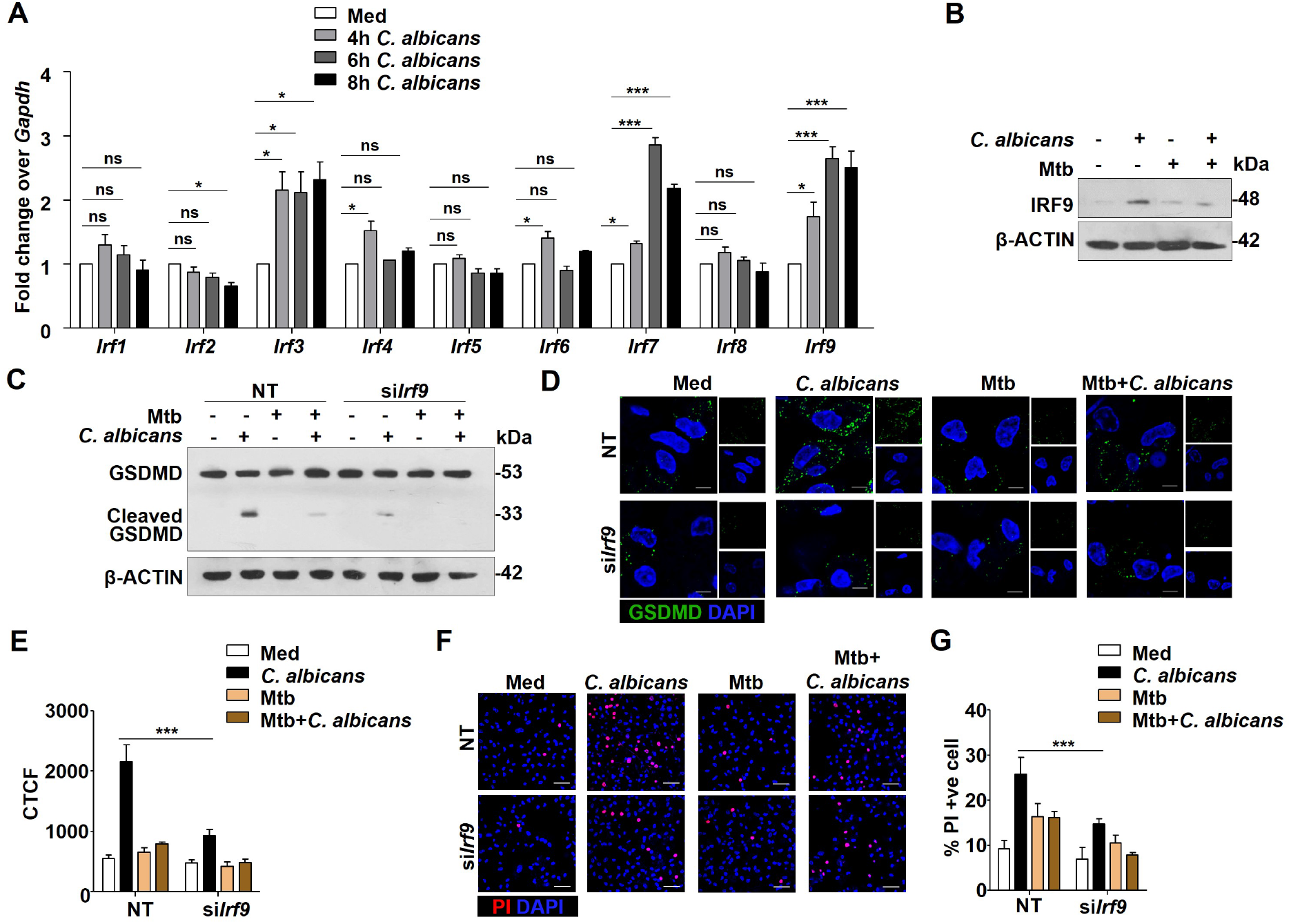
IRF9 mediates *C. albicans* induced pyroptosis. (A) Transcript level of members of IRF family genes upon infection with *C. albicans* to peritoneal macrophages for indicated time. (B) Immunoblot assessment for IRF9 in peritoneal macrophages infected with *C. albicans* for 6 h or Mtb for 18 h or co-infection with Mtb for 12 h followed by *C. albicans* for 6 h. (C-G) Peritoneal macrophages transfected with siRNA specific to *Irf9* or NT siRNA were infected with *C. albicans* for 6 h or Mtb for 18 h or co-infected with Mtb for 12 h followed by *C. albicans* for 6 h. (C) Level of cleaved GSDMD, (D) representative IF images of GSDMD and (E) its quantification. (F) Representative images of PI and DAPI stained peritoneal macrophages and (G) its quantification. Error bar represents the mean ± SEM for at least 3 independent experiments; ns, not significant; *, p < 0.05; **, p < 0.01; ***, p < 0.001 (One-way ANOVA followed by Tukey’s test for panel A, two-way ANOVA followed by Tukey’s test for panel E and G). Blots are representative of 3 independent experiments. Scale bar, 5 μm(D) and 40 μm(F); original magnifications 63X (D) and 40X (F); Med, medium; CTCF, corrected total cell fluorescence; NT, non-targeting.

Furthermore, prior Mtb infection suppressed the *C. albicans* induced expression of IRF9 at protein level without any alteration at the transcript level (Fig 3B and S3A). siRNA mediated knock down of IRF9 during *C. albicans* infection supressed the cleavage as well as surface localization of GSDMD (Fig 3C-E) with augmented viability of macrophages (Fig 3F-G). Altogether, these results suggest that IRF9 is essential for execution of pyroptosis during *C. albicans* infection.

### COP1 targets IRF9 for proteasomal degradation during co-infection

To elucidate the mechanism responsible for regulating the expression of IRF9 at protein level but not at transcript level during co-infection, we assessed for proteasomal degradation of IRF9. It has been reported that IRF9 undergoes proteasomal degradation during Varicella zoster virus infection [30]. We observed that treatment with proteasome inhibitor (MG132) in macrophages rendered Mtb incapable of suppressing *C. albicans* driven expression of IRF9 (Fig 4A). Further, to identify the E3 ubiquitin ligase which is responsible for proteasomal degradation of IRF9, we used a bioinformatics platform named UbiBrowser (http://ubibrowser.ncpsb.org) and as shown in Fig 4B, top 20 predicted E3 ubiquitin ligases that can target IRF9 with a confidence score of more than 0.660 were identified. Among computationally predicted ligases, genome wide expression analysis of human TB patients sample demonstrated upregulation of four E3 ubiquitin ligases namely PML, RFWD2, TOPORS and TRIM25 (Fig. 4C) [31]. Previously, a study from our lab suggested that *M. bovis* BCG induces expression of COP1 (*Rfwd2*), a RING finger family ubiquitin ligase [32].

**Fig. 4.**
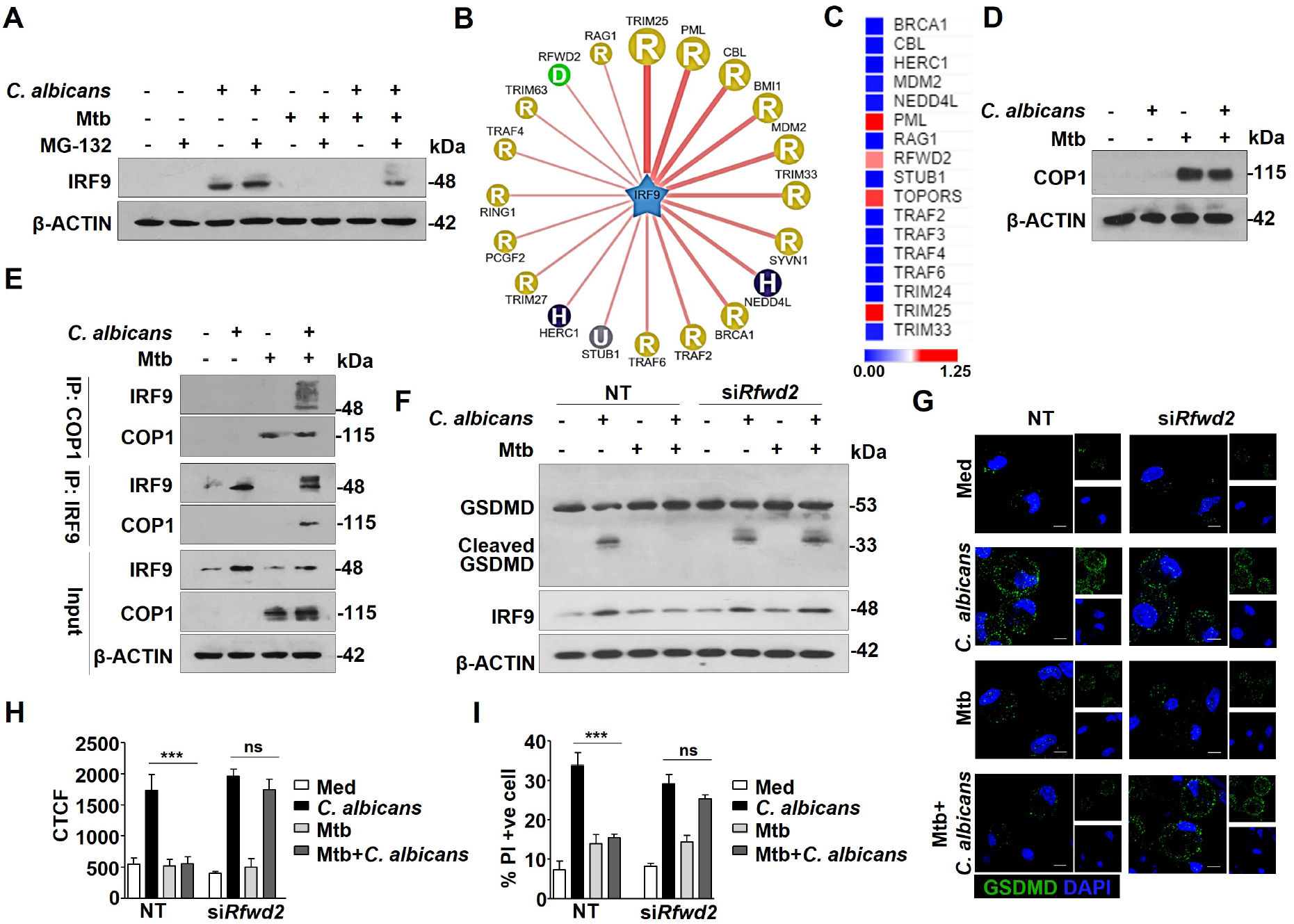
COP1 targets IRF9 for proteasomal degradation during co-infection. (A) Peritoneal macrophages were infected with Mtb for 18 h or *C. albicans* for 6 h or co-infected with Mtb for 12 h followed by *C. albicans* infection for 6h. (A) Protein level of IRF9 was assessed with or without treatment with MG132 for 4 h prior to completion of experiment. (B) Bioinformatics prediction indicating potential E3 ubiquitin ligases for IRF9. (C) Heat map depicting the transcript levels of E3 ubiquitin ligase in Tuberculosis patients. Peritoneal macrophages were infected with *C. albicans* for 6 h or Mtb for 18 h or co-infected with Mtb for 12 h followed by *C. albicans* infection for 6h. (D) Expression of COP1 at protein level. (E) Immunoprecipitation for IRF9 or COP1 followed by immunoblotting for IRF9 and COP1. (F-I) NT or *Rfwd2* siRNA transfected peritoneal macrophages were infected *C. albicans* for 6 h or Mtb for 18 h or co-infected with Mtb and *C. albicans*. (F) Cleaved GSDMD and IRF9 protein levels, (G) representative IF images of GSDMD and (H) its quantification, (I) quantification of PI positive macrophage to assess viability. Error bar represents the mean ± SEM for at least 3 independent experiments; ns, not significant; ***, p < 0.001 (One-way ANOVA followed by Tukey’s test), each blot is representative of 3 independent experiments. Scale bar, 5 μm; original magnifications 63X (G); Med, medium; CTCF, corrected total cell fluorescence; NT, non-targeting.

Therefore, we assessed for COP1 expression and observed a significant induction upon Mtb infection while *C. albicans* infection did not induce COP1 expression. Level of COP1 during co-infection remained unaltered compared to Mtb infection alone (Fig 4D, Fig S3B). As predicted, COP1 and IRF9 were found to be interacting during co-infection (Fig 4E) and pulldown experiments showed ubiquitinated form of IRF9 and treatment with MG132 rescued degradation of IRF9 (Fig S3C). To further establish the role of COP1 in modulating IRF9 level, siRNA mediated knockdown of COP1 was carried out and results demonstrated similar levels of IRF9 in *C. albicans* and co-infected macrophages (Fig 4F), indicating that Mtb induced COP1 targets IRF9 for proteasomal degradation. Moreover, knock down of COP1 during co-infection did not suppress *C. albicans* induced pyroptosis as evident from the level of cleaved GSDMD and its surface localisation (Figure 4F-H). An increase in macrophage death was observed upon knock-down of COP1 during co-infection as compared to cells transfected with non-targeting siRNA (Fig 4I). Altogether, these results suggest that during co-infection, Mtb driven COP1 directly marks IRF9 with K-48 linked poly-ubiquitination which targets IRF9 for proteasomal degradation and suppresses pyroptosis induced by host to clear *C. albicans* infection.

### *M. tuberculosis* activated PKCζ-WNT pathway suppresses pyroptosis by regulating COP1

Having established the role of COP1 in degradation of IRF9, we attempted to delineate signalling pathway involved in Mtb induced COP1 expression. We and others have shown that Mtb infection could trigger WNT/β-Catenin signalling pathway in various cellular contexts and also promoter of COP1 demonstrates several binding sites for β-Catenin, an effector of WNT signalling cascade [7,33]. siRNA mediated knockdown of β-Catenin or pharmacological abrogation of WNT signalling compromised the ability of Mtb to induce the expression of COP1 (Fig 5A, S4A and S4B). In line with these result, enhanced recruitment of β-Catenin at the promoter of COP1 was observed upon Mtb infection (Fig 5B) suggesting the crucial role for WNT/β-Catenin signalling in regulating the expression of COP1. Furthermore, PKCζ (Protein Kinase C ζ) is known to positively regulate WNT/β-Catenin pathway. PKCζ-mediated phosphorylation stabilises Dishevelled3 (an integral part of WNT signalling) and positively regulates the nuclear localisation of β-Catenin [34,35]. Additionally, it has been previously reported that PKC family of kinases are activated upon mycobacterial infection [36,37]. To this end, we utilized PKCζ inhibitor along with the dominant negative form of PKCζ and observed that PKCζ is required for the activation of WNT pathway during Mtb infection (Fig 5C and S4C). As mentioned before, WNT pathway regulates the expression of COP1 and accordingly, Mtb induced COP1 expression was found to be dependent on PKCζ (Fig 5D and S4D). Collectively, PKCζ-WNT/β-Catenin signaling axis contributes to enhance expression of COP1 during Mtb infection. Further, to assess the role for WNT pathway in regulation of IRF9 and pyroptosis during co-infection, siRNA mediated knock down of β-Catenin was performed. In consonance with the results of COP1 knockdown, silencing of β-Catenin also showed comparable levels of IRF9 and cleavage of GSDMD in co-infected macrophages to that of *C. albicans* alone infected macrophages (Fig 5E-F). These evidences suggest that silencing of β-Catenin abrogates the expression of COP1 and rescues IRF9 from proteasomal degradation during co-infection.

**Fig 5.**
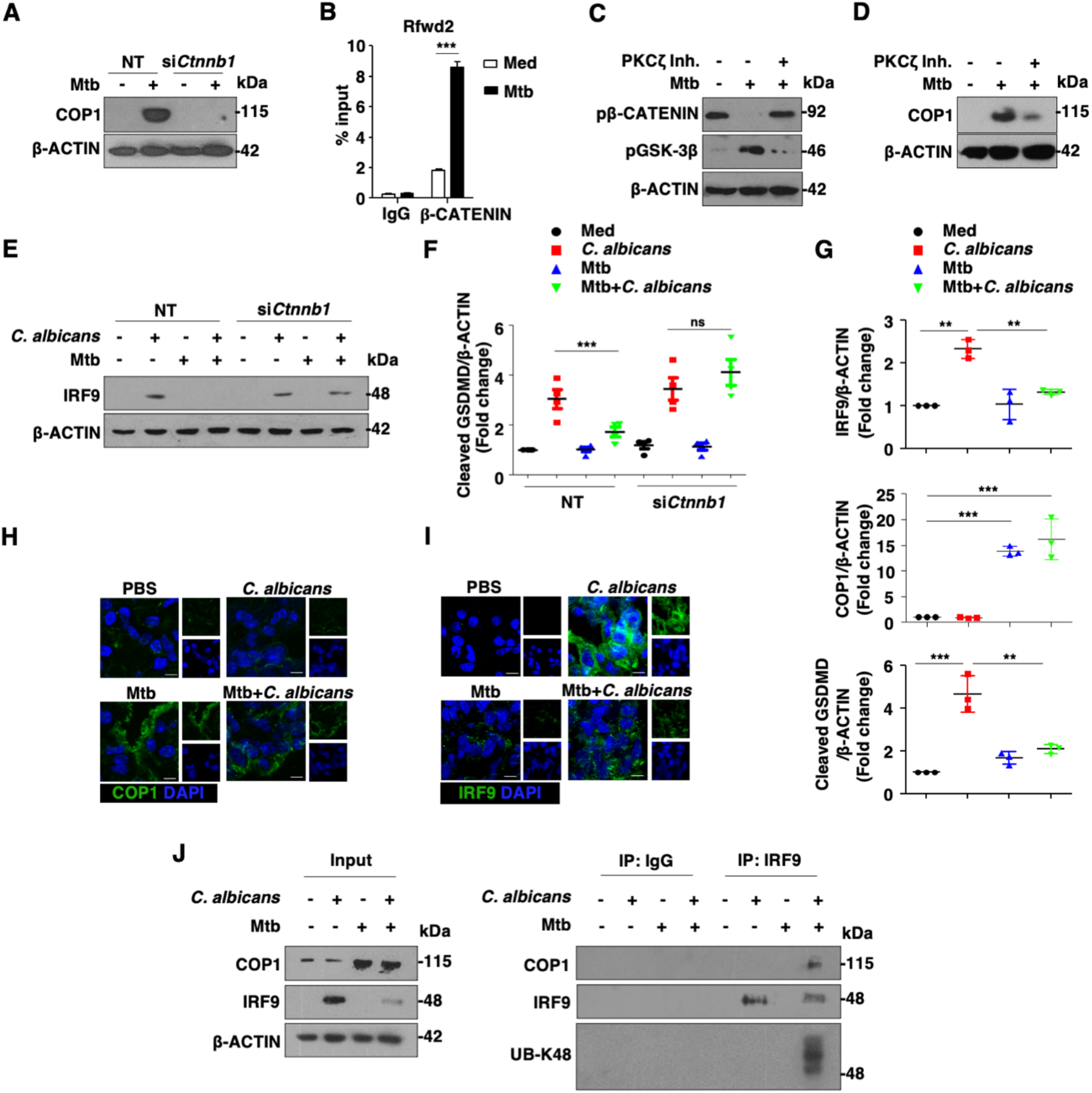
*M. tuberculosis* activated PKCζ-WNT pathway suppresses pyroptosis by regulating COP1. (A) Expression of COP1 was assessed in mouse peritoneal macrophages which were transfected with *Ctnnb1* siRNA or NT siRNA followed by 12 h infection with Mtb. (B) ChIP analysis for β-CATENIN recruitment at the promoter of *Rfwd2* in murine peritoneal macrophages infected with Mtb for 12 h. (C) Peritoneal macrophage were treated with PKCζ inhibitor 1h prior to Mtb infection for 1h and the whole cell lysate was assessed for pβ-CATENIN and pGSK3β expression by western blotting. (D) COP1 expression in peritoneal macrophage treated with PKCζ inhibitor followed by 12 h of Mtb. (E-F) Mouse peritoneal macrophages were transfected with *Ctnnb1* or NT siRNA followed by infection with Mtb for 18 h or *C. albicans* for 6 h or co-infection with Mtb or *C. albicans*, (E) western blot for IRF9 and (F) densitometry analysis of cleaved GSDMD. (G-J) Mice were infected with *C. albicans* or Mtb or co-infected with *C. albicans* and Mtb, (G) densitometry analysis for the protein level of IRF9, COP1 and cleaved GSDMD in lung lysate, representative IF images of (H) COP1 and (I) IRF9 in the lung cryosections. (J) Immunoprecipitation of IRF9 followed by immunoblotting for COP1, Ub-K48 and IRF9 in lung lysate of mice infected with *C. albicans* or Mtb or co-infected with *C. albicans* and Mtb. Each Blot is representative of 3 independent experiments. Scale bar, 5 μm; original magnifications 63X; Error bar represents the mean ± SEM for at least 3 independent experiments; ***, p < 0.001; ns, not significant (Student’s t-test for panel B, One-way ANOVA followed by Tukey’s test for panel F and G); NT, non-targeting; Inh., Inhibitor

To examine the *in vivo* relevance of mechanism elucidated during co-infection, levels of COP1, IRF9 and cleaved GSDMD were assessed in the lung tissues of Mtb infected, *C. albicans* infected and co-infected mice. In line with data derived from *in vitro* studies, COP1 was found to be induced upon Mtb infection as well as during co-infection. However, *C. albicans* induced expression of IRF9 was impaired during co-infection and pyroptosis was suppressed as evident from GSDMD cleavage (Fig 5G-I). Also, *in vivo* co-immunoprecipitation assay suggested that IRF9 interacts with COP1 and gets ubiquitinated in mice coinfected with Mtb and *C. albicans*. Together, these results depicts the intricate cross-talk between the two pathogens in regulating IRF9 and COP1 levels both *in vitro* and *in vivo*.

## Discussion

In general, pyroptosis protects the host against infection and helps in clearing the pathogen such as *Salmonella*, *Listeria*, *Shigella* etc. [38–40]. On the contrary, exorbitant inflammation and pyroptosis can be detrimental for the host as reported for inflammatory bowel disease, myocardial infarction, neurodegenerative diseases etc. [40–43]. Release of inflammatory cytokines as well as cell lysis during pyroptosis requires many downstream events including Caspase-1 dependent pore formation. Moreover, Caspase-1 deficient mice are more susceptible to *Francisella* and *Listeria* infection [44,45]. These observations clearly indicate that pathogen and host compete with each other to regulate pyroptosis, balance of which governs the survival of the host. Surprisingly, though pyroptosis is well reported during *C. albicans* infections [14,15], its relevance during *in vivo* infection remains obscure.

In this context, we found that inhibition of Caspase-1 leads to suppression of pyroptotic cell death thus leading to excessive *C. albicans* burden and associated mice mortality. However, we would like to mention possible role for impaired Th1 responses in Caspase-1 deficient mice thus advocating roles for other processes [46]. Moreover, our results demonstrates that inhibition of Caspase-1 lead to mice mortality within 3 days post *C. albicans* infection suggesting role for innate immune responses as the major contributor at such early time. Further, IL-1β and IL-18 which are released during pyroptosis, leads to recruitment of immune cell and hence innate control of *C. albicans* infection.

Reports suggests that type I interferons (IFN) play central role in inflammasome activation and pyroptosis during various infection scenarios ranging from bacterial, viral to fungal infection [47–49]. Further, type I IFN genes are known to be regulated by members of IRF family [50]. In this regard, we report a pivotal role of IRF9 in enhancing pyroptosis during *C. albicans* infection.

As previously mentioned, a number of clinical studies report the co-existence of *C. albicans* with *M. tuberculosis* in human patients [2–5]. Despite this, extensive studies on cross-talk between these two pathogens and host responses is limited. In view of these observations, we have established a mouse model to study Mtb-*C. albicans* co-infection and we found a significant decrease in survival of mice upon co-infection corroborated by high *C. albicans* load. Mice survival studies and *C. albicans* CFU assays suggest that pyroptosis protects host during *C. albicans* infection and Mtb co-infection suppress pyroptosis hence leading to severe pathogenesis and increased mortality. In addition, mouse model established in this study can further be utilized as a therapeutic model to study role of various small molecule inhibitors during coinfection.

E3 ubiquitin ligases serve regulatory functions during inflammatory disease and hence may serve as a potential drug target[51]. In regards to Mtb infection, a number of E3 ubiquitin ligases are reported to play central role in Mtb pathogenesis. Mtb infection in Smurf1^−/−^(SMAD Specific E3 Ubiquitin Protein Ligase 1) mice shows exacerbated lung inflammation with increased Mtb burden [52]. NEDD4 E3 ubiquitin protein ligase enhances autophagy and subsequent Mtb killing [53].

In the current study, we found induced expression of COP1 by virulent strain of Mtb both *in vitro* and *in vivo* corroborating earlier finding from our lab wherein vaccine strain of mycobacterium (*M. bovis* BCG) was reported to induce expression of COP1 [32]. Regulation of COP1 by both vaccine and virulent strain prompted us to speculate the involvement of PRR (pathogen recognition receptor)-driven signalling. Further, TLR2 is the major PRR established to get activated during Mtb infection [54,55]. Moreover, we found that PKCζ regulates the expression of COP1 which in turn is known to physically interacts with TLR2 leading to activation of downstream signalling [36,56]. Interestingly, studies attribute activation of WNT signalling by both Mtb as well as *C. albicans* infection [33,57]. However, in the current study, WNT-responsive expression of COP1 was observed only with Mtb infection but not during *C. albicans* infection alone. WNT pathway can cross talk with other signalling pathways such as HIPPO, MAPK and Rac-Rho [58–60]. Therefore, it leaves us with an open question whether co-factors or signalling cross talk with active WNT pathway is different during Mtb and *C. albicans* infection.

Altogether, our study shows alluring cross talk during *C. albicans* and Mtb co-infection revealing a significant role for pyroptosis as a regulator for manifesting a successful *C. albicans*-Mtb co-infection (Fig 6).

**Fig 6.**
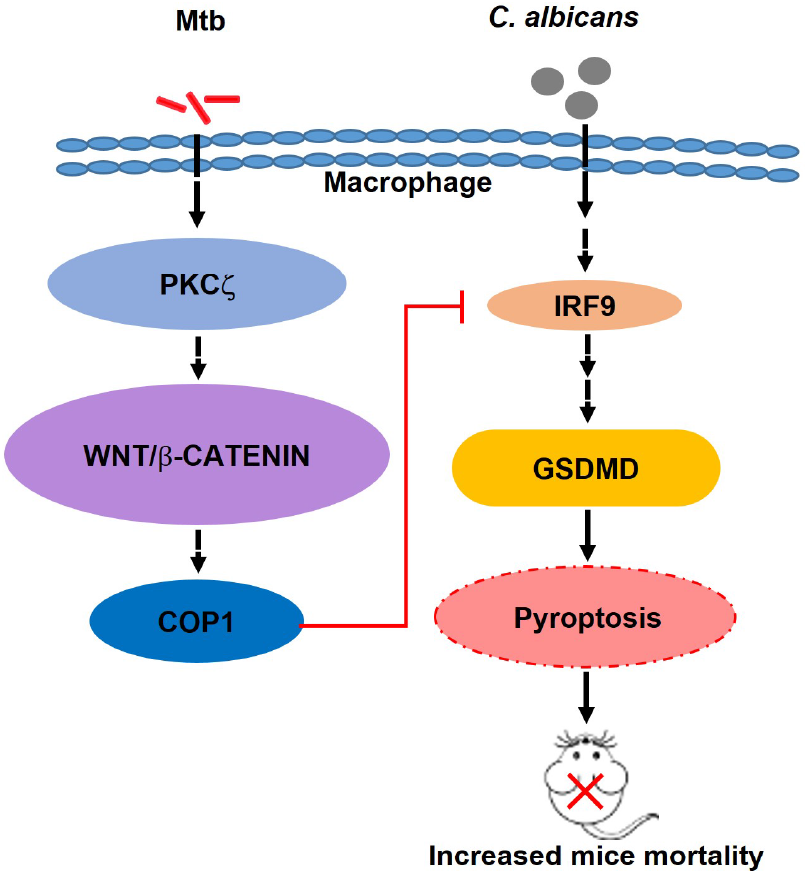
Schematic: PKCζ-WNT/β-Catenin signaling driven expression of COP1 upon Mtb infection leads to proteasomal degradation of *C. albicans*-triggered expression of IRF9 and thus causes inhibition of pyroptosis.

## Acknowledgements

We thank Central Animal facility at IISc for maintaining and providing mice for experimentation. Members of K.N. Balaji Laboratory are acknowledged for timely help during the course of this investigation. PKCζ DN construct was obtained as kind research gifts from Dr. Jae-Won Soh, Inha University, Korea. We acknowledge BSL-3 facility and Dr. RS Rajmani for permitting our i*n vitro* and *in vivo* experiments with Mtb H37Rv. We thank Dr. Ravi Manjithaya for allowing us to use slide scanner microscopy facility at JNCASR, Bangalore. Anusha Jettur, Navya and Puneeth of the MCBL confocal facility are acknowledged for their generous help. Dr. Madhavi Naik, Consultant Pathologist at St. Theresa Hospital, Bangalore is thanked for assistance in analysing the pulmonary pathology of mice. This work was supported by funds from the Department of Biotechnology (DBT No. BT/PR27352/BRB/10/1639/2017, DT.30/8/2018 and BT/PR13522/COE/34/27/2015, DT.22/8/2017 to K.N.B) and the Department of Science and Technology (DST, EMR/2014/000875, DT.4/12/15 to K.N.B.), New Delhi, India. K.N.B. thanks Science and Engineering Research Board (SERB), DST, for the award of J. C. Bose National fellowship (No.SB/S2/JCB-025/2016, DT.25/7/15) and for the funding (SP/DSTO-19-0176, DT.06/02/2020). The authors thank DST-FIST, UGC Centre for Advanced Study and DBT-IISc Partnership Program (Phase-II at IISc BT/PR27952/INF/22/212/2018) for the funding and infrastructure support. Fellowships were received from IISc (PP, GKL) and UGC (BB).

## Author contributions

BB contributed in conceptualization, investigation, formal analysis, manuscript draft preparation and editing. PP contributed in conceptualization, investigation, formal analysis, manuscript draft editing; GKL contributed in investigation, manuscript draft editing; KNB contributed in conceptualization, formal analysis, supervision, manuscript review and editing.

## Materials & Correspondence

Dr. Kithiganahalli Narayanaswamy Balaji, Department of Microbiology and Cell Biology, Indian Institute of Science, Bangalore 560012, Karnataka, India. Phone: +91-80-22933223, Fax: +91-80-23602697, E-mail address.

## Supplementary Figures and legends

**Fig. S1.**
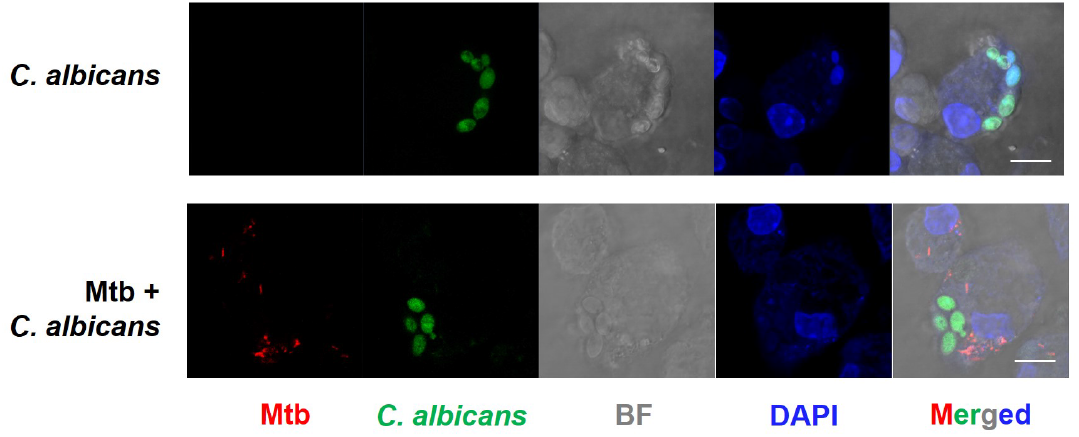
Uptake of *C. albicans* in macrophage. Representative IF image of DAPI stained peritoneal macrophages infected with *C. albicans* (CEC2684) for 2 h or Mtb for 14 h or co-infected with Mtb for 12 h followed by *C. albicans* infection for 2 h. Scale bar, 10 μm; original magnifications 100X.

**Fig. S2.**
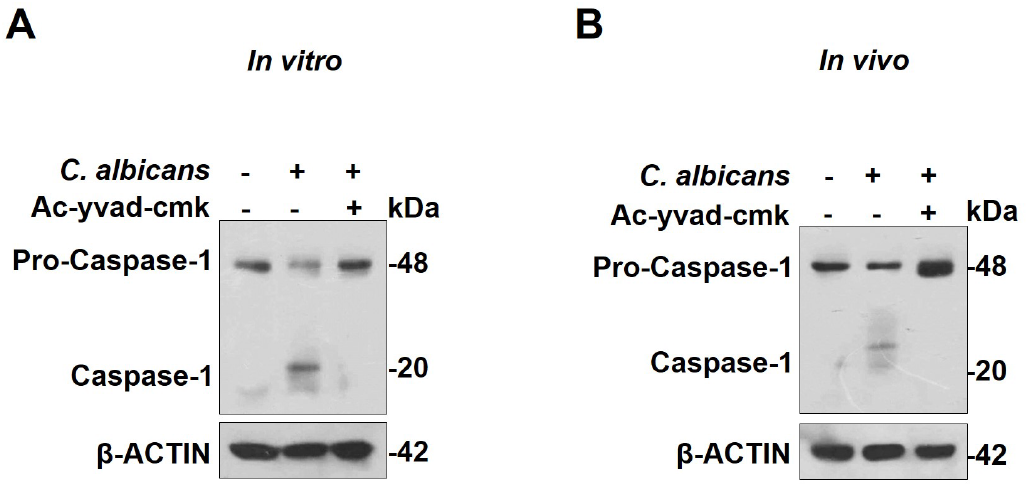
Ac-yvad-cmk inhibits *C. albicans* induced Caspase-1 activity. Protein level of total and cleaved Caspase-1, (A) in murine peritoneal macrophages infected with *C. albicans* for 6 h with and without 1 h prior treatment with Ac-yvad-cmk and (B) in lung lysate of BALB/c mice harvested at 24 h post *C. albicans* infection with or without treatment of Ac-yvad-cmk. Blots are representative of 3 independent experiments.

**Fig. S3.**
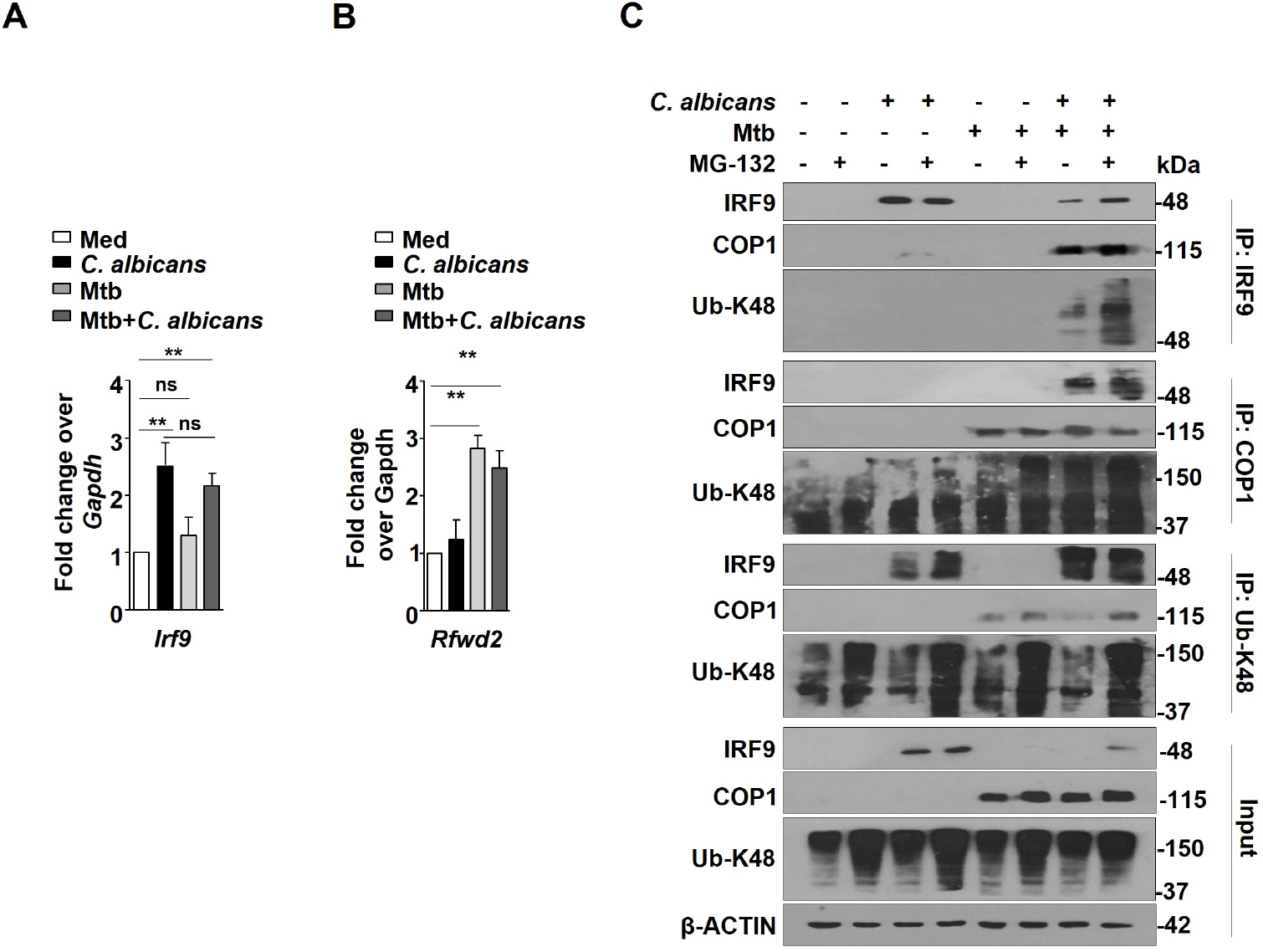
COP1 mediates proteasomal degradation of IRF9. Peritoneal macrophages were infected with *C. albicans* for 6 h or Mtb for 18 h or co-infected with Mtb for 12 h followed by *C. albicans* infection for 6 h. Transcript level of *Irf9* (A) and *Rfwd2* (B) were assessed. (C) Pull down of IRF9, COP1 or Ub-K48 was performed in presence or absence of MG-132 treatment given 4 h prior to completion of experiment followed by immunoblotting for indicated protein. Each blot is representative of 3 independent experiments. Error bar represents the mean ± SEM for at least 3 independent experiments; ns, not significant; **, p < 0.01 (One-way ANOVA followed by Tukey’s test).

**Fig. S4.**
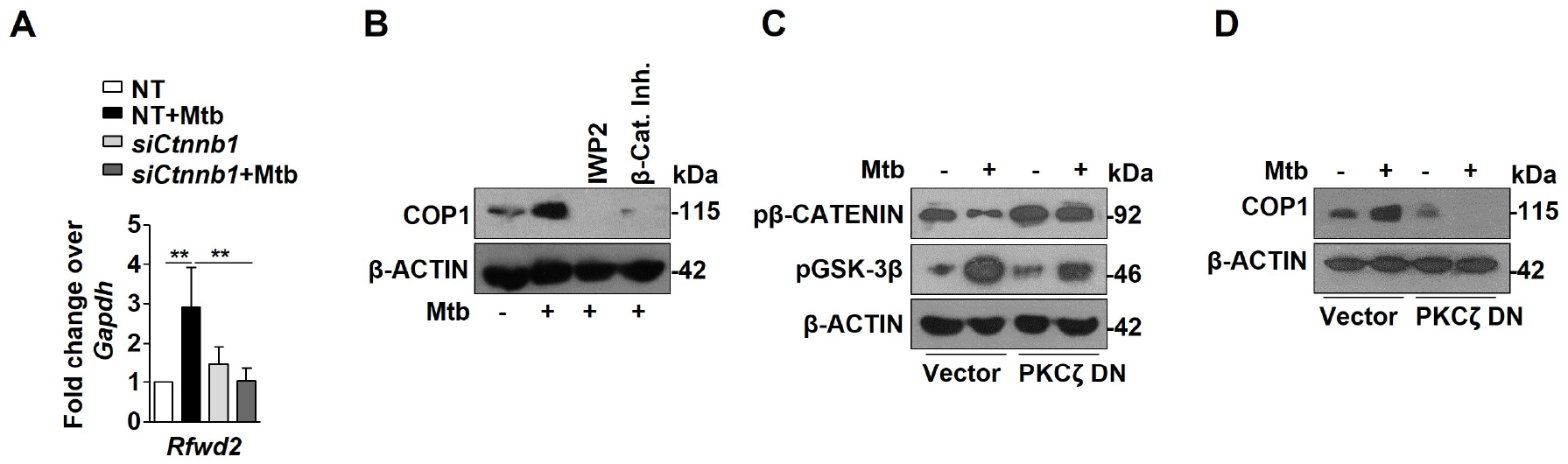
PKCζ activated WNT signaling axis enhances COP1 expression. (A) Transcript level of *Rfwd2* were assessed in murine peritoneal macrophages transfected NT or *Ctnnb1* siRNA post 12 h Mtb infection. (B) Expression of COP1 by western blotting in mouse peritoneal macrophages pre-treated with IWP2 and β-Catenin inhibitor for 1 h followed by Mtb infection for 12 h. RAW264.7 macrophages transfected with empty vector or PKCζ DN construct followed by Mtb infection for 1 h (C) or 12h (D) were assessed for indicated proteins. All data represents the mean + SEM from 3 independent experiments, **, p < 0.01 (One-way ANOVA followed by Tukey’s test) and all blots are representative of 3 independent experiments. DN, dominant negative; Inh., inhibitor; NT, non-targeting.

**Supplementary Table 1:**
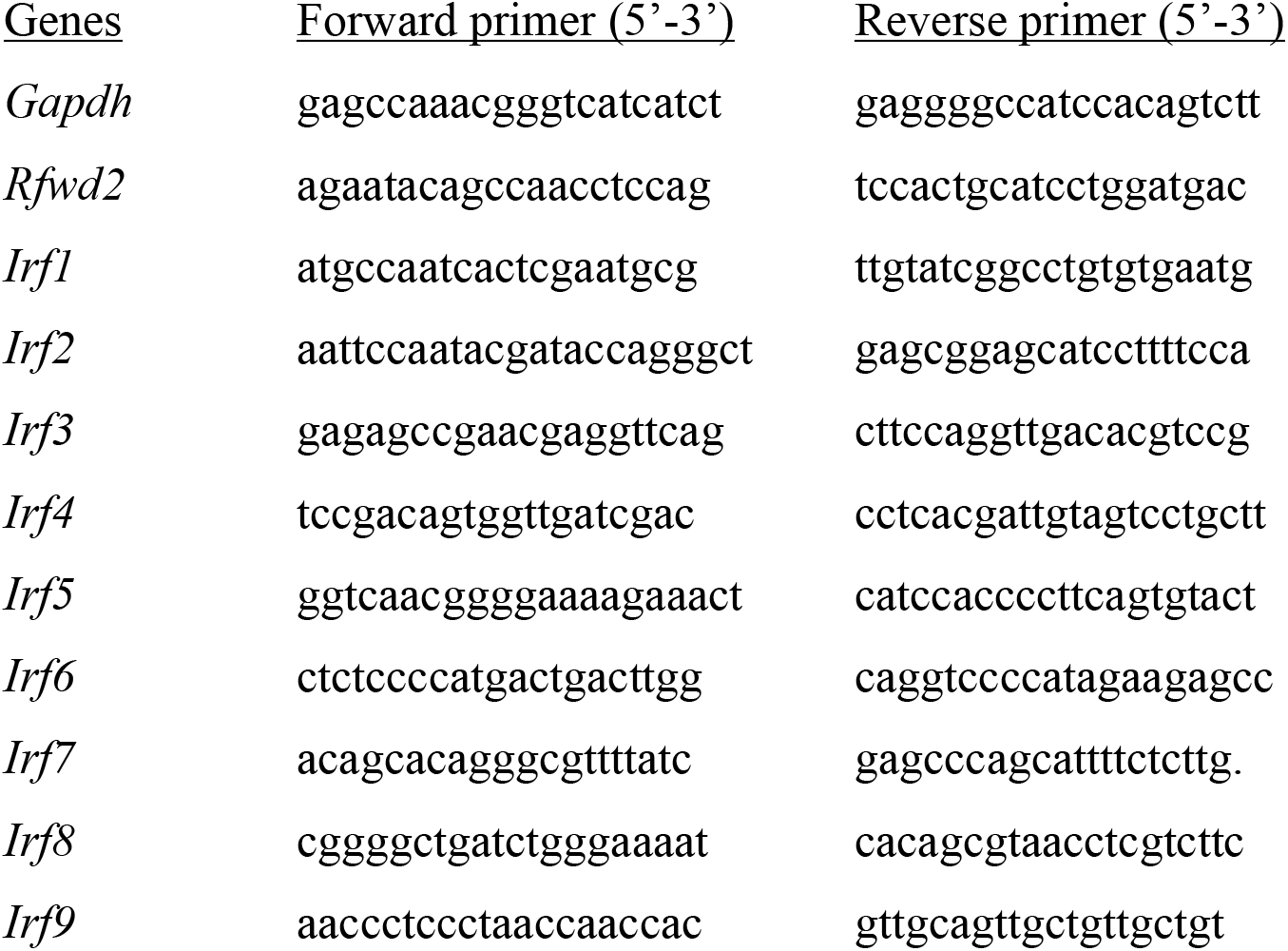
List of primers for qRT-PCR.

**Supplementary Table 2:**
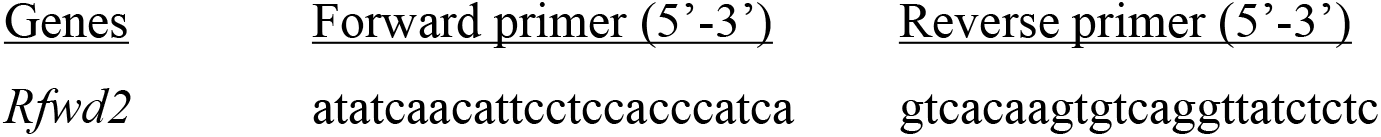
List of primers for ChIP assay.

